# The Nuclear Pore Complex acts as a hub for pri-miRNA transcription and processing in plants

**DOI:** 10.1101/2024.10.24.620027

**Authors:** Lucia Gonzalo, Delfina Gagliardi, Camila Zlauvinen, Tomasz Gulanicz, Agustín L. Arce, Josefina Fernandez, Damian A. Cambiagno, Agnieszka Zienkiewicz, Artur Jarmolowski, Pablo A. Manavella

**Affiliations:** Instituto de Agrobiotecnología del Litoral, CONICET, Universidad Nacional del Litoral, Santa Fe, Argentina; Instituto de Hortofruticultura Subtropical y Mediterránea “La Mayora” (IHSM “La Mayora”), Universidad de Málaga-Consejo Superior de Investigaciones Cientificas (UMA-CSIC), Campus Teatinos, 29010 Málaga, Spain; Centre for Modern Interdisciplinary Technologies, Nicolaus Copernicus University in Toruń, 87-100 Toruń, Poland; Cátedra de Bioinformática, Facultad de Bioquímica y Ciencias Biológicas, Universidad Nacional del Litoral, 3000 Santa Fe, Argentina; Department of Gene Expression, Institute of Molecular Biology and Biotechnology, Adam Mickiewicz University, Poznan, Poland; Current address: Unidad de Estudios Agropecuarios (UDEA), Instituto Nacional de Tecnología Agropecuaria (INTA) - Consejo Nacional de Investigaciones Científicas y Técnicas (CONICET), Córdoba, Argentina

## Abstract

The regulation of miRNA biogenesis and movement is essential for plant development and environmental responses. HASTY (HST), a karyopherin protein, has been implicated in miRNA biogenesis and movement, though its role in non-cell-autonomous miRNA movement remains unclear. Through a genetic screen, we identified that mutations in the *HAWAIIAN SKIRT (HWS)* gene suppress the developmental defects of *hst* mutants by restoring miRNA movement. Our findings show that HWS interacts with nuclear transport factors and nuclear pore complex (NPC) components, including NUP1, positioning HWS as a regulator of miRNA nuclear export. Using microscopy and fluorescence *in situ* hybridization, we showed that pri-miRNA transcription, and likely their co-transcriptional processing, occurs at the nuclear pore. Notably, we uncovered an antagonistic relationship between HST and HWS in regulating *MIRNA* transcription at the NPC and AGO1 loading, which could explain the observed changes in miRNA movement. HST promotes the association of *MIRNA* loci with the NPC, likely facilitating co-transcriptional processing, while HWS negatively regulates this process by targeting MEDIATOR complex subunits to degradation detaching, consequently, the processing complex from the NPC. Furthermore, our data provides evidence of spatial coordination of miRNA transcription, biogenesis, and movement highlighting a novel role for the NPC in the miRNA pathway.

## Introduction

HST was initially identified as a factor controlling the transition from the juvenile phase to the adult phase and flowering, two critical stages of development in which miRNAs play a crucial role (1). Mutant plants deficient in HST exhibit an accelerated transition between these phases, resulting in abnormal leaf growth, defective leaf arrangement, and sterility (1,2). This protein belongs to the karyopherin family, a group of proteins required for the export and import of biological molecules through the nuclear pore complex. HST’s closest homolog in animals is EXPORTIN 5 (XPO5), which is involved in shuttling miRNA precursors from the nucleus to the cytoplasm (2–9). This homology, coupled with the fact that miRNAs levels are disturbed in HST mutants, led researchers to assume that HST’s function in the miRNA pathway was related to the export of miRNAs to the cytoplasm (2,7). However, subsequent studies refuted this assumption by demonstrating that although HST is capable of shuttling between the nucleus and the cytoplasm, it is dispensable for miRNA nuclear export (10).

A recent study indicates that HST is involved in regulating the non-cell autonomous functions of miRNAs by controlling their cell-to-cell movement, though the underlying mechanism remains unclear (11). Interestingly, it was shown that HST modulates miRNA biogenesis by acting as a scaffold between DICER-Like 1 (DCL1) and the MEDIATOR complex, enabling the recruitment of the processing complex to *MIRNA* loci (10), a process that ultimately favors the co-transcriptional processing of pri-miRNAs (12). In plants, miRNAs can be loaded into AGO1 in the nucleolus and exported as an active RISC complex (13). However, it has been postulated that once loaded into AGO1, miRNAs lose their inter-cell mobility (14), making the pool of free cytoplasmic miRNAs (15) crucial for the non-cell autonomous miRNA pathway. Curiously, most mobile miRNAs show a high rate of co-transcriptional processing, a process that, like miRNA movement, is tightly controlled by HST (12). This creates an intriguing network where AGO1 loading, co-transcriptional processing, and HST appear to interact to define mobile miRNAs. Adding another layer of complexity, both the cytoskeleton and the nuclear pore complex (NPC) play roles in AGO1 loading and thus in miRNA movement (16,17). In animals, TREX2 (Transcription Export Complex 2) and THO-TREX tether genes to the NPC during transcription and coordinate Pol II transcription with mRNA export to the cytoplasm (18–21). In plants, mutants in core components of TREX-2, such as THP1 and SAC3A, show defects in miRNA biogenesis and nuclear export (17). Additionally, both THP1 and SAC3A physically interact with Pol II and the key *MIRNA* transcription regulators CELL DIVISION CYCLE 5 (CDC5) and NOT2B. They also interact with SERRATE (SE) and C-TERMINAL DOMAIN PHOSPHATASE-LIKE 1 (CPL1), which promotes HYL1 dephosphorylation (17,22–24). Strikingly, THP1 also interacts with NUP1 in the nuclear envelope. In *thp1* and *nup1* mutants, the nuclear export of both miRNAs and AGO1 is reduced, suggesting that TREX-2 promotes the export of miRISC complexes (17). The THO/TREX complex, another known complex associated with the NPC, shows similar involvement in miRNA processing. Mutants in THO2, a core protein of THO/TREX complex, exhibit severe defects in miRNA biogenesis, although no direct interaction with known miRNA biogenesis components has been detected. However, it has been shown that THO2 interacts with miRNA precursors, facilitating their association with the miRNA processing complex (25). The combination of evidence suggests that the nuclear pore and its accessory proteins are likely cross-talking with the miRNA processing complex in plants.

While NPCs are surrounded by regions of uncondensed chromatin, the nuclear envelope is closely associated with heterochromatin (26). TPR (Translocated Promoter Region), a component of the nuclear basket that forms filaments, modulates gene expression by excluding heterochromatin from regions near the NPC and promoting uncondensed chromatin associated with active transcription (27). In Arabidopsis, a homolog of TPR known as NUCLEAR PORE ANCHOR (NUA) has been identified (28). This protein is part of the NPC and regulates gene expression, DNA spatial organization, and mRNA export. Previous studies have linked the absence of this protein to decreased FLC transcripts (4); recently, it has been shown that transcription of FLC occurs associated with the nuclear pore, enhancing its expression and facilitating an efficient transport of the resulting mRNA to the cytoplasm (29).

Here, we provide compelling evidence that miRNA transcription, and likely pri-miRNA processing, occurs associated to the nuclear pore in Arabidopsis. We demonstrate that HST, likely utilizing its exportin-like properties, actively recruits *MIRNA* loci to the nuclear pore vicinity, creating a spatially optimized environment for miRNA maturation and export. Conversely, HWS appears to antagonize this process by inducing the degradation of scaffold proteins and detaching the miRNA processing complex from the NPC. This suggests an antagonistic regulatory mechanism in which HST and HWS modulate the accessibility of miRNA loci to the nuclear pore. These dynamics offer new insights into miRNA nuclear export and their role as mobile signaling molecules. The interplay between HST and HWS highlights an unexpected aspect of miRNA biogenesis that allows us to consolidate previous reports into a comprehensive mechanism of miRNA production where spatially optimized transcription and processing may be the key aspect.

## Results

### Mutations in the gene encoding HWS suppress the phenotypes of *hst-15* mutants

We performed a suppressor screen on *hst-15* mutants to further explore the biological function of HST in Arabidopsis using an unbiased genetic approach. *hst-15* seeds were mutagenized with ethylmethane sulfonate (EMS), and second-generation seedlings were screened for suppression of the typical *hst-15* leaf phenotype, primarily characterized by long hyponastic leaves. Approximately 20,000 mutagenized seeds were screened, and 18 individuals from 11 M1 seed pools were identified with varying degrees of phenotype suppression (Figure S1A).

To identify the gene responsible for suppressing the *hst-15* phenotype, we backcrossed the obtained mutants with the *hst-15* parental line and used a mapping-by-sequencing approach to search for causal mutations (30). In the first batch of sequenced plants, we identified SNPs in coding regions associated with the reversion of the phenotype in seven mutant plants (Supplementary Table 1). Among the identified SNPs, we found a substitution resulting in a Proline to Serine non-synonymous mutation in the gene encoding IMPA2. The isolation of this mutant serves as a proof of concept for the utility of the screen given the previously established association of IMPA2 with HST function (10).

Among the sequenced mutants, we identified the causal mutation for mutant 35-23 in a small region on chromosome 3. Within this region, we found a typical EMS-induced C>T (G>A) substitution that results in a glycine to arginine mutation at position 238 (G238R) in the gene encoding HAWAIIAN SKIRT (HWS) (AT3G61590) (Figure 1A). HWS is an F-BOX protein that interacts with classic components of the SCF complex (31,32). *HWS* mutants display fused sepals, a trait associated with a defective CUC1 miR164 regulatory node (33,34). Interestingly, *hws* mutations suppress phenotypes caused by miRNA target mimicry and were associated with target-induced microRNA degradation in Arabidopsis (35,36). Moreover, in an interesting genetic interaction, mutations in proteins associated with the miRNA pathway, including HST, can revert the *hws* phenotype (37).

**Figure 1.**
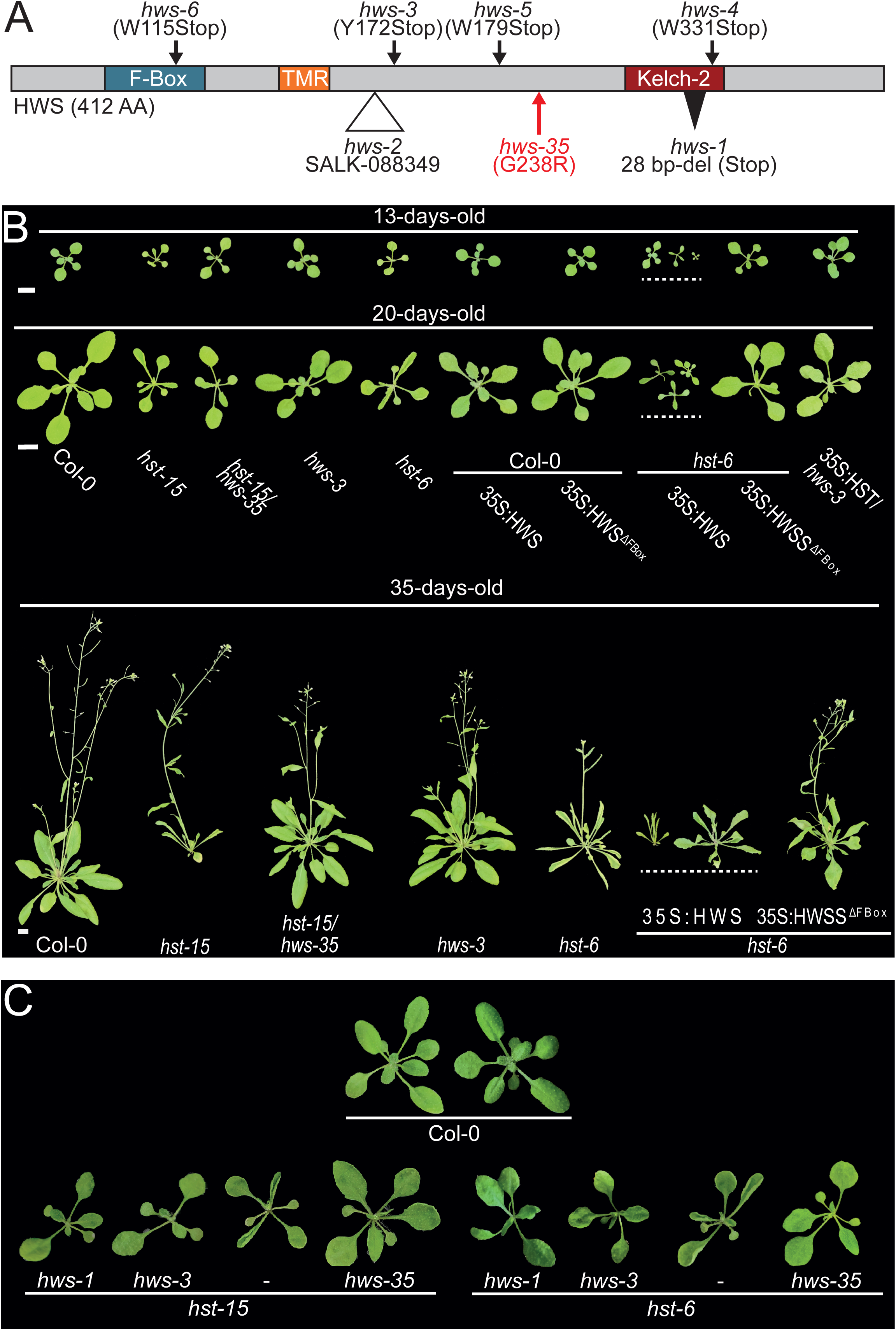
A mutation in *HWS* reverses the *hst* phenotype. A-Schematic representation of HWS. Previously published mutant alleles of *HWS* are indicated in black. The new *HWS* mutant allele identified in this study (*hws-35*) is marked in red. Single nucleotide mutations are displayed with arrows, T-DNA insertions with empty triangles, and deletions with black triangles. The blue box represents the F-Box domain, the orange box the TMR domain, and the dark red box the Kelch-2 domain. B-Phenotypes of 13-, 20-, and 35-day-old Col-0 wild type and mutant plants. Scale bar= 1 cm. C-Phenotype of 20-day-old *HST/HWS* double-mutants combining different alleles of both mutants. In B and C, plants were imaged individually and mounted in a single black background square to facilitate comparison.

The isolated mutant, named *hws-35* hereafter, showed partial suppression of the *hst-15* phenotype, most notably the loss of leaf hyponasty (Figure 1B, S1A). Overexpression of *HWS* in the double mutant background restored and even exacerbated the *hst-15* phenotype, confirming that the reversion of the *hst-15* phenotype was caused by the mutation in *hws* (Figure S1B). This exacerbation of the *hst* mutant phenotype was also observed when HWS was overexpressed in the *hst-6* mutant allele but not when transformed into a wild-type background (Figure 1B). This suggests that HWS may target an HST compensatory element, exacerbating the phenotype only when HST is absent. Interestingly, overexpression of a mutant version of HWS lacking its F-BOX domain (HWS^ΔFBox^) only partially reverted the *hst-15* phenotype and did not exacerbate the *hst-6* phenotype, indicating the relevance of this domain in phenotype suppression (Figure 1B, S1B). To further confirm that a mutation in *HWS* is sufficient to suppress the *hst-15* phenotype, we crossed this mutant, and *hst-6*, with other *hws* alleles (*hws-1* and *hws-3*) and observed that they also suppressed the hyponastic leaf phenotype of *hst* mutants further confirming the genetic interaction between both genes (Figure 1C).

### Suppression of the *hst-15* phenotype in *hws* mutants is not explained by impaired HST turnover

HWS is an F-BOX-domain-containing protein predicted to be part of the SCF complex, which regulates protein turnover. Then, if HWS targets HST, the absence of active HWS could increase HST levels, potentially explaining the suppressed phenotype of the double mutant. This scenario would only be plausible if *hst-15* is a hypomorphic allele that still produces a partially active protein. When genotyping *hst-15* mutants, we were unable to obtain amplification products using primers around the T-DNA insertion site at exon 18 (Figure 2A and S2, red lines). In fact, we were only able to amplify the T-DNA-containing region in *hst-15* when using primers located far from the putative insertion site (Figure 2A and S2, black line), and even in this case, the amplification fragment was much smaller than the wild-type version (Figure S2). When performing the mapping by sequencing of our suppressor screen candidates we re-sequenced *hst-15* mutants, we thus used this data to re-assemble the *HST* locus to detect the T-DNA insertion. Surprisingly, we found that not only was the T-DNA insertion missing in the mutant, but a large region of the *HST* gene, encompassing exons 16 to 19, was also deleted in *hst-15* (Figure 2B). This deletion explains the absence of PCR amplification in *hst-15* when using T-DNA flanking primers and is likely a consequence of natural T-DNA removal, which resulted in the partial loss of the flanking regions.

**Figure 2.**
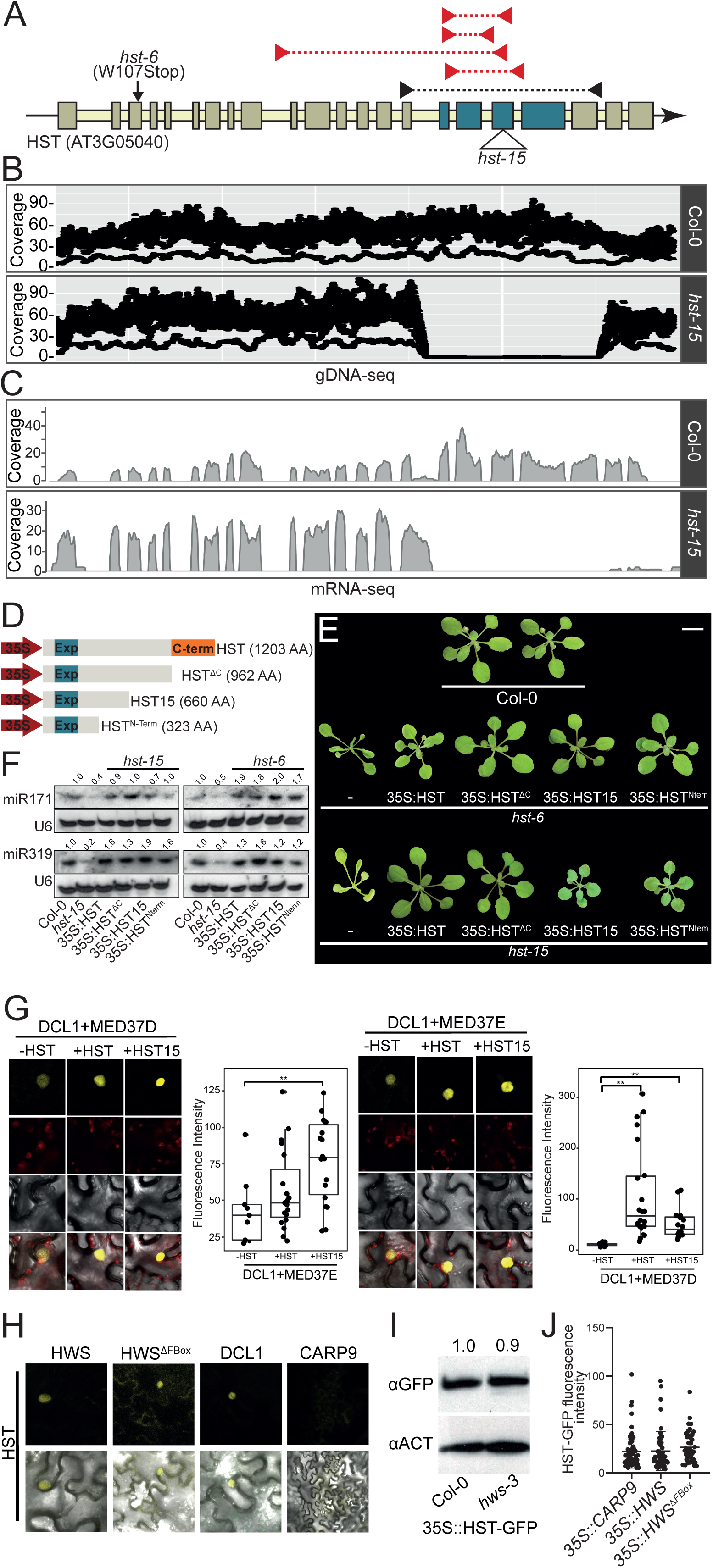
*hst-15* is a hypomorphic allele. A-Schematic representation of the *HST* gene. Rectangles represent exons, while the thick line refers to introns. Blue rectangles indicate deleted exons due to T-DNA loss. The empty triangle marks the position where the T-DNA was annotated for the *hst-15* allele, and the *hst-6* allele is marked with a black arrow. Red triangles connected by dashed lines represent the non-amplified fragments in *hst-15*; black triangles connected by a dotted line represent the obtained amplification product in this allele. B and C-Coverage of genomic DNA sequencing (B) and mRNA sequencing (C) reads over the *HST* locus in wild-type and *hst-15* mutant plants. D-Schematic representation of the different versions of *HST* used for phenotype complementation: HST: full version; HST^ΔC^: version without the C-terminal domain; HST15: truncated version resulting from T-DNA removal in the *hst-15* mutant; HST^N-term^: version only including the N-terminal region of the protein. The exportin domain is shown in blue. Length of each construct is noted in brackets. E-Phenotype of 20-day-old *hst-15* and *hst-6* mutants complemented with different versions of HST. F-RNA blots for detection of mature miR171 and miR319 in the complemented *hst* plants. U6 was used as a loading control. Signal intensity was calculated with ImageJ and normalized to U6. Ratios of signal intensities of mutants to control are noted above each gel. G-TriFC assays showing the interaction between DCL1-MED37D and DCL1-MED37E in *N. benthamiana* leaves in the presence or absence of HST and HST15 versions, respectively. Fluorescence intensity was quantified with ImageJ and expressed relative to background. Error bars represent SEM, and p-values below 0.05 (*) or 0.01 (**), in an unpaired t-test, were considered significant. H-Interaction between HST-HWS and HST-HWS^ΔFBox^ measured by BiFC. DCL1 was a positive control for HST interaction, and CARP9 was a negative control. I-Western blot detecting HST:GFP levels in transgenic wild-type and *hws-3* mutant plants. ACTIN was quantified as a loading control. J-HST-GFP levels, as measured by fluorescent intensity in nuclei of plants expressing HST under its native promoter and co-expressing HWS, HWS^ΔFBox^, or CARP9, as a negative control, under a 35S promoter.

mRNA-seq data from *hst-15* revealed that this genomic deletion led to a shorter transcript that retained part of intron 15, containing a stop codon 21 amino acids after the end of exon 15 (Figure 2C). This transcript would translate into a truncated protein of 660 amino acids, approximately half the size of the full-length HST protein.

This particularity raises the possibility that the truncated HST-15 protein, if deregulated in *hws-35,* could be sufficient to partially complement the *hst* mutant phenotype. To test this hypothesis, we created three truncated versions of HST: one missing the C-terminal domain (HST^ΔC^), one mimicking the truncated version present in *hst-15* (HST15), and a smaller version containing only the N-terminal domain (HST^N-term^) (Figure 2D). We then overexpressed these constructs in *hst-15* and *hst-6* mutants and assessed phenotype complementation. All constructs, with varying levels of penetrance, were able to partially complement the phenotypes caused by *hst* mutation, particularly the characteristic hyponastic leaves and selected miRNA levels (Figure 2E and F). The truncated HST15 protein was even sufficient to promote the interaction between DCL1 and MED37D/MED37E (Figure 2G), a known HST scaffold activity (10).

While this evidence suggests that the hypomorphic activity of the truncated protein in *hst-15* could explain the phenotype reversion if overaccumulated in *hws-35,* other data suggest otherwise. First, we observed that *hws* mutations not only reverted the *hst-15* phenotype but also the *hst-6* phenotype, which, given its premature stop codon, is likely a null allele (Figure 1D). Furthermore, although we detected HWS-HST interaction via BiFC (Figure 2H), we did not observe any change in HST protein levels in *hws-3* mutants (Figure 2I) nor when overexpressing HWS or HWS^ΔFBox^ (Figure 2J). Altogether, our data reveal the hypomorphic nature of the *hst-15* allele and put a cautionary note on this allele for further studies. However, our findings also provide evidence that the suppressor effect of *hws* mutations on *hst-15* is unlikely to be due to defective HST turnover.

### Mutations in *HWS* suppress the *hst-15* phenotype by restoring miRNA movement

The phenotypic characteristics of *hst* mutants have been attributed to impaired cell-to-cell movement of miRNAs, particularly miR390, which reduces the non-cell-autonomous functions of miRNAs. This disruption impairs the formation of a spatial gradient of mobile sRNAs required for leaf polarity, leading to the observed leaf developmental defects (11). Therefore, it is possible that the restoration of miRNA movement in the double mutants leads to the observed suppression of the phenotype.

In agreement with previous reports (11), basic fuchsin staining of the protoxylem and metaxylem in 5-day-old *hst-6* and *hst-15* roots showed signs of impaired miR165/166 non-cell-autonomous activity from the endodermis to the stele, resulting in xylem defects. These defects are characterized by the presence of only one protoxylem file (instead of the normal pair), which was occasionally interrupted by gaps (Figure 3A). None of these defects were observed in *HWS* mutants. Furthermore, the root defects detected in *hst* mutants disappeared in *hws-35/hst-15* double mutants (Figure 3A). Confirming the connection between HWS and HST functions, similar root-architecture defects were observed in plants overexpressing *HWS,* but not in those overexpressing *HWS^ΔFBox^*(Figure 3A). These results indicate that HWS, likely through its F-BOX domain, influences HST function in controlling non-cell-autonomous miRNA activity, which in turn affects plant development.

**Figure 3.**
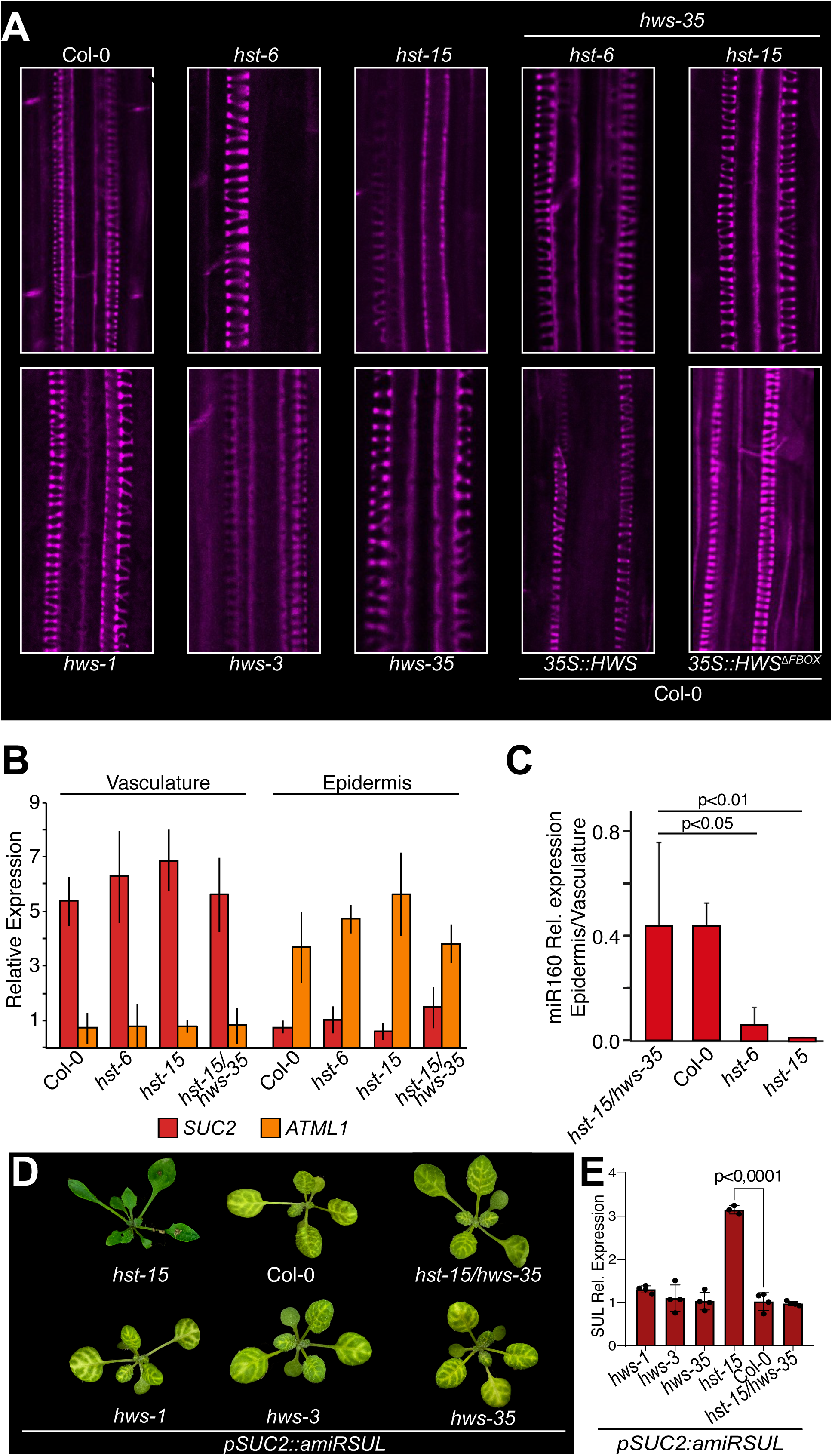
Impaired non-cell-autonomous miRNA movement in *hst* mutants is restored by *HWS* mutation. A-Basic fuchsin staining of 5-day-old roots showing the protoxylem and metaxylem. *hst-15*, *hst-6*, and HWS overexpression lines show root defects, while the double *hst/hws* mutant restores normal root morphology. B-RT-qPCRs confirming proper separation of epidermis (*ATML1*) and vasculature (*SUC2*) tissues. C-Ratio of miR160 levels between epidermis and vasculature as measured by RT-qPCR. *hst* mutants show defective miRNA movement from vasculature to epidermis, while the double *hst/hws* mutant restores movement to wild-type levels. Error bars represent SEM. D-amiRSUL-silencing chlorosis phenotype in *pSUC2::amiRSUL* transgenic *Arabidopsis* plants in different mutant backgrounds. E-RT-qPCR of *SUL* mRNA, showing higher expression in *hst-15,* where movement is not impaired, compared to other genotypes. Error bars represent SEM.

To further confirm this effect on cell-to-cell miRNA movement, we analyzed the levels of miR160 in Meselect separated leaf vasculature and epidermis (38,39). The transcription of *MIR160* is restricted to the leaf vasculature, but mature miR160 moves from its biogenesis source in the vasculature to the leaf epidermis to regulate its target genes. This allows us to quantify miR160 cell-to-cell movement by separating the vasculature and epidermis (39). To confirm the purity of the tissue separation, we measured *SUC2* and *ATML1* expression in the isolated tissues, as these genes are exclusively expressed in the vasculature and epidermis, respectively (Figure 3B). As previously observed, miR160 movement was impaired in both *hst-15* and *hst-6* mutants, a defect that was completely reversed in the double *hst15/hws-35* suppression lines (Figure 3C).

We also tested miRNA movement using the well-established *pSUC2::amiRSUL* transgenic system (40). In this system, an artificial miRNA (amiRSUL), which targets the magnesium chelatase subunit CHLORINA42 mRNA (SUL), is specifically expressed in phloem companion cells (CC) under the control of the *SUC2* promoter (*pSUC2::amiRSUL*). In this system, miRNA movement is scored by the extension of the chlorosis area proximal to the veins, which is caused by the movement of amiRSUL from CCs (11,40) . As previously reported, *hst* mutations reduce amiRSUL movement, as indicated by both a reduced area of chlorosis and elevated levels of *SUL* mRNA (Figure 3D and E). Conversely, although the *HWS* mutation alone does not affect amiRSUL movement, it is capable of restoring amiRSUL movement in *hst-15* mutants (Figure 3D and E). This further suggests that the absence of *HWS* can restore the impaired miRNA movement observed in *hst* mutants. It has been suggested that non-cell-autonomous miRNA function rely on the pool of AGO1-unloaded miRNAs (11,14,16). Although the efficiency of miRNA loading into the global cellular pool of AGO1 remains largely unchanged in the *hst* mutant (11), we observed an impaired miRNA loading into AGO1 in *hws* mutants as measured by RIP-qPCR experiments (Figure S3). This inefficient AGO1 loading may allow specific miRNAs to move out of the cell explaining the restoration of the non-cell-autonomous functions we observed in the double mutants.

### HWS interacts with the nuclear pore and most exportin/importin proteins in Arabidopsis

To explore the crosstalk between HST and HWS, we analyzed mass-spectrometry data for both proteins (10,35) and searched for shared interactors. We identified over 110 proteins shared between the two datasets (Figure 4A), including IMPA2, RAN1, and MED37, which we previously found necessary for HST nuclear/cytoplasmic shuttling and its recruitment to miRNA loci (10) (Supplementary Table 2). HWS also interacted with several miRNA-related factors, such as AGO1, AGO2, CRM1, and SE. We validated some of these interactions using an alternative approach, confirming HWS interactions with SE, IMPA2, and RAN1 (Figure S4A and S4B). A GO term (Biological Processes) analysis of the shared interactors revealed a notorious enrichment for the term “intracellular protein transport” (Figure 4B).

**Figure 4.**
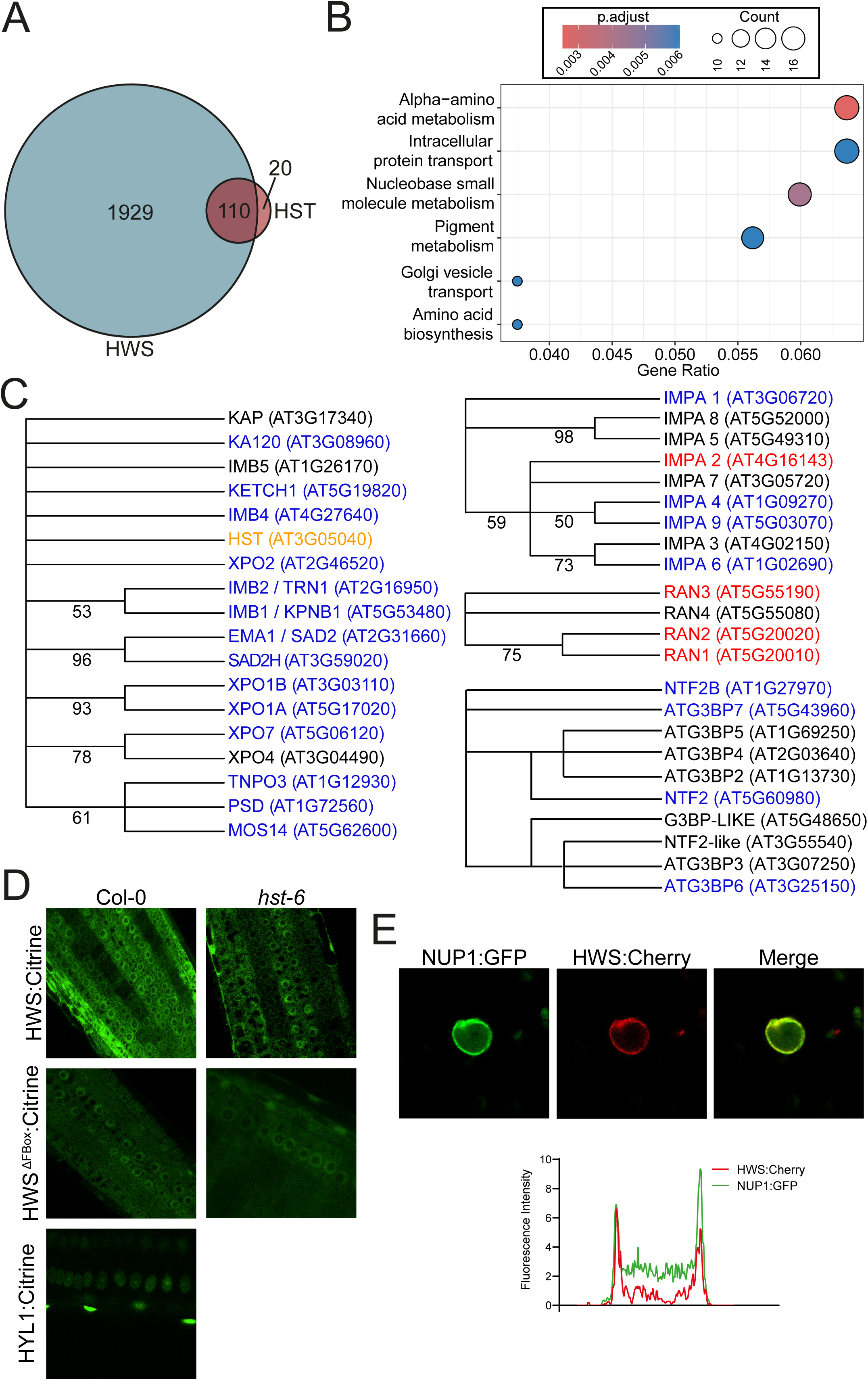
HWS interacts with the Nuclear Pore Complex (NPC). A-Venn diagram showing proteins interacting with both HWS and HST as detected by mass spectrometry. B-Gene ontology analysis showing enrichment of proteins based on the GO term Biological Processes, after filtering for log_2_ FC > 10. Protein counts are represented by circle size, and p-values are shown in a color gradient. C-Phylogenetics of Alpha Exportins, Beta Importins, RAN GTPases, and WIP proteins. Proteins interacting with HWS (blue) or both HWS and HST (red) as detected by mass spectrometry are marked. HWS-HST interaction (orange) was detected by alternative approaches. D-Localization of HWS:Citrine and HWS^ΔFBox^:Citrine in wild-type and *hst-6* roots. HYL1:GFP was used as a nuclear localization control. E-Co-localization of pNUP1:NUP1:GFP with 35S:HWS:Cherry in *N. benthamiana.* Co-localization was quantified by ImageJ as the overlap between the red and green signals in a transection of the cells and plotted as a histogram.

Interestingly, the HWS-mass spectrometry analysis also revealed interactions with a large portion of the exportin, importin, nuclear transport factors, and RAN GTPases identified in Arabidopsis (Figure 4C). Out of the 41 nuclear export/import-related proteins, HWS appeared to interact with 27 (84% of all exportins) (Figure 4C). Given this overwhelming overrepresentation, it is likely that HWS interacts with the remaining proteins under specific cellular conditions or in certain cell types, as the mass spectrometry did not detect the HWS-HST interaction we observed by other approaches (Figures 2H).

The overrepresentation of karyopherins in the HWS interactome suggests that HWS may regulate nuclear/cytoplasmic shuttling. Therefore, the suppression of *hst-15* phenotype by *hws* mutations could involve miRNA intracellular movement. Although previous evidence ruled out a role for HST in miRNA nuclear export (10,13,17,41) and confirmed that HST’s modulation of non-cell-autonomous miRNA functions is independent of miRNA nuclear/cytoplasmic shuttling (11), we decided to evaluate whether HWS affects subcellular miRNA movement.

To test this hypothesis, we fractionated Arabidopsis nucleus and cytoplasm and measured the relative accumulation of mature miRNAs in different mutant and overexpressing lines. As previously reported, *hst* mutants did not show a general altered miRNA partitioning (Figure S4C). This normal distribution was also observed in the *HWS* overexpressing lines, but an apparent nuclear miRNA retention was observed for *hws* mutants (Figure S4C). The modulation of the subcellular trafficking of several miRNA-related proteins is biologically relevant. For example, HYL1 turnover depends on its nucleus/cytoplasm distribution, controlled by the karyopherin KETCH1 (22,23,42,43). Similarly, AGO1 nuclear import is critical for miRNA loading, while its shuttling to the cytoplasm is required for target mRNA silencing (13). To assess whether HWS affects the subcellular distribution of miRNA-related proteins, we co-transformed fluorescent-tagged versions of HYL1, HST, AGO1, CARP9, RAN1, and IMPA2 together with HWS or HWS^ΔFBox^ and quantified the fluorescence intensity ratio between the nucleus and the cytoplasm. No statistically significant differences in subcellular distribution were observed for most proteins beside HST where there was an apparent nuclear retention in those cells overexpressing HWS but not HWS^ΔFBox^ (Figure S4D).

Alternatively, the overrepresentation of karyopherins in the HWS interactome may indicate that HWS acts at a common node for all these importins and exportins. Notably, the strongest signal detected in the HWS mass spectrometry experiment was from NUCLEAR-PORE ANCHOR (NUA - AT1G79280), a well-characterized component of the nuclear pore basket and a docking point for karyopherins on their way through the nuclear pore complex. Although with lower intensity, we also identified NUP93 as a HWS interactor in the mass spectrometry experiment. NUP1, another component of the nuclear pore basket, has also been linked to miRNA activity in previous studies (17). Given the large size of NUA (237-kDa), and the difficulty in working with the entire protein (44), we performed many of the following experiments using NUP1 as a marker for the nuclear pore basket, as it is more suitable for the design of molecular experiments. Despite also having a broader nuclear distribution, we detected HWS localization on the nuclear membrane using confocal microscopy (Figure 4D). This localization was previously observed but overlooked in an earlier work (Figure S6 of (35)). In line with HWS’s interaction with the NPC, we found that the peripheral localization of HWS coincided with NUP1 in co-localization experiments (Figure 4E). BiFC experiments, using a tagged version of a NUP1 genomic construct, confirmed the interaction between HWS and NUP1 at the nuclear pore basket observed as interaction signal forming a nuclear peripheral ring (Figure S5).

The observation of HWS interacting with the nuclear pore basket raises the possibility that the suppression of the *hst-15* mutant phenotype in *hws-35* mutants is due to a more permissive nuclear pore, allowing miRNAs or miRNA-related proteins to move more efficiently through the nuclear membrane, even in the absence of HST. However, our results indicate that this does not appear to be the case (Figure S4).

### HST and HWS Antagonistically Define *MIRNA* Transcription at the Nuclear Pore

Beyond its putative function as an exportin, we previously showed that HST participates in the miRNA pathway as a scaffold, recruiting the miRNA processing complex to *MIRNA* loci by interacting with the MEDIATOR complex (10). This recruitment promotes co-transcriptional processing of nascent pri-miRNAs, a process that we found to be impaired in *hst* mutants (12). To quickly measure impaired co-transcriptional processing of pri-miRNAs, we quantified the levels of nascent unprocessed pri-miRNAs associated with *MIRNA* loci after chromatin IP. As previously reported, we observed a significant reduction in co-transcriptional pri-miRNA-processing in *hst-15* mutants, a defect that was largely rescued in the double *hst-15/hws-35* plants (Figure 5A). Interestingly, we observed that *hws-35* mutants exhibited increased co-transcriptional miRNA processing compared to wild-type plants (Figure 5A), suggesting that while miRNA processing is not directly affected by HWS (35), it negatively regulates co-transcriptional miRNA processing.

**Figure 5.**
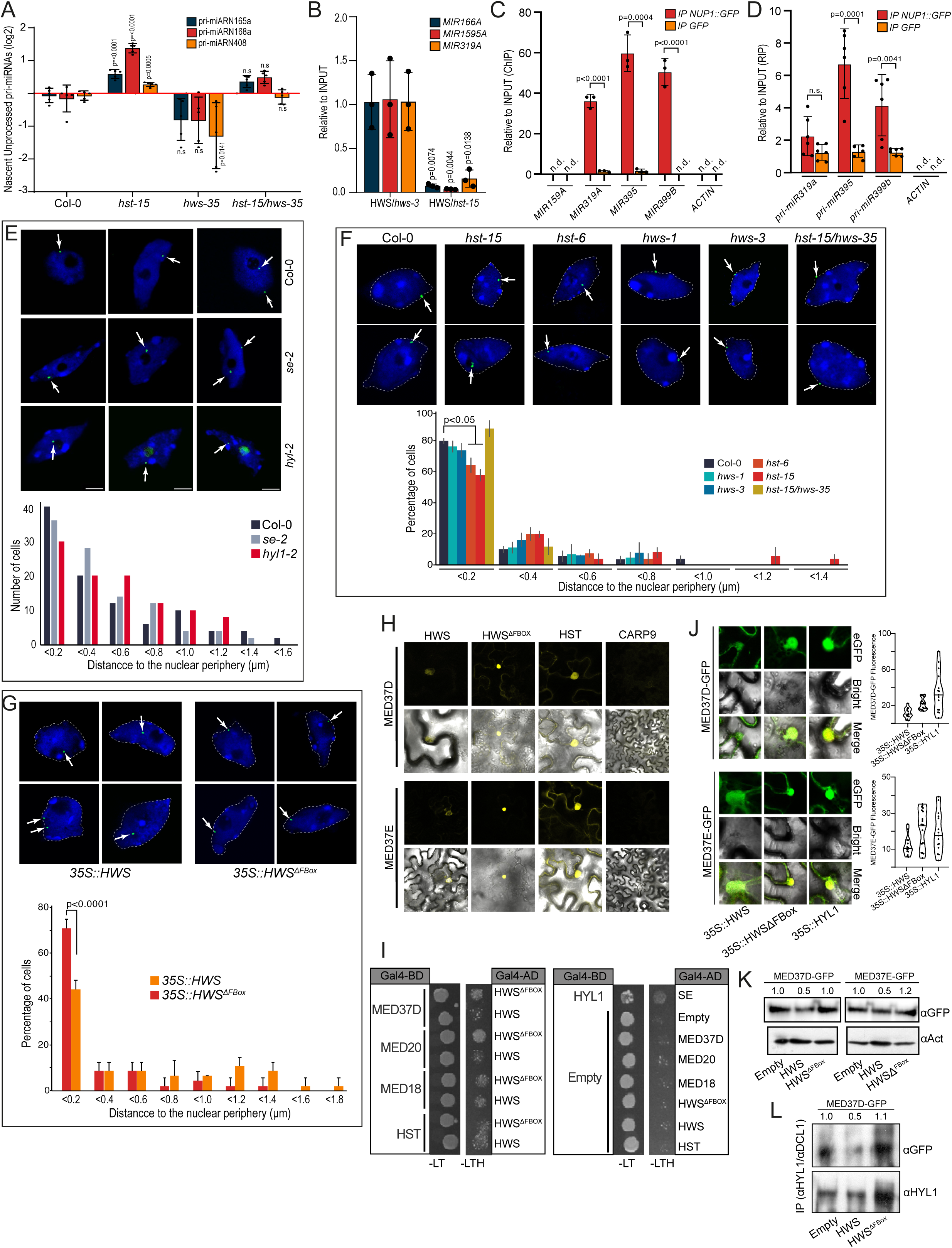
HST and HWS antagonistically regulate *MIRNA* transcription at the NPC. A-Quantification of unprocessed pri-miRNAs associated with chromatin in different genotypes measured by RIP-qPCR. B-ChIP-qPCR analysis of *MIRNA* loci associated with HWS in *hws-3* and *hst-15* mutants expressing *HWS-Citrine*, showing interaction of HWS with *MIRNA* loci and its dependence on HST. Results are expressed relative to *hws-3*/*35S::HWS*-Citrine plants. C and D-ChIP-qPCR (C) and RIP-qPCR (D) assays in *N. benthamiana* transformed with pNUP1::NUP1-GFP, detecting *MIRNA* loci and pri-miRNAs interaction with the NPC. Non-transformed leaves were used as a negative control. Non detected signal is noted as n.d. (A-D) qPCR values are from n=3 biologically independent samples, presented as mean ± SEM. P-values were calculated in a two-tailed, unpaired, t-test. E, F, and G-Visualization of the *MIR156A* locus in the nucleus of wild-type, mutant plants (E and F), and plants overexpressing HWS or HWS^ΔFBox^ (G) using fluorescence in situ hybridization (FISH). The blue signal resulting from DAPI staining was used to delimitate the nucleus envelope, which is marked with discontinues grey lines. *MIRNA* locus is observed in green, and its distribution quantified as the average percentage of cells with the *MIR156A* signal at an indicated distance, measured with ImageJ, from the nuclear peripheral zone. Scale bars, 2 *µ*m. H-BiFC assay of HWS and HWS^ΔFBox^ with MED37D and MED37E. HST was used as a positive interaction control, and CARP9 as a negative control. I-Yeast two-hybrid assay of HWS and HWS^ΔFBox^ with different MEDIATOR subunits and HST. SE - HYL1 interaction was the positive control, and empty GAL4-AD and GAL4-BD constructs served as negative controls. J-Quantification of 35S::MED37D-GFP and 35S:MED37E:GFP fluorescence intensity in the presence of 35S::HWS or 35S::HWS^ΔFBox^ to assess their degradation. HYL1-GFP was used as a negative control. Representative images are paired with dot plot graphs showing quantifications. K and L-Quantification of MED37D-GFP and MED37E-GFP protein levels in plants stably transformed with 35S::HWS or 35S::HWS^ΔFBox^ either in the total protein fraction (K) or in a protein fraction obtained after HYL1 / DCL1immunoprecipitation (L). ACTIN and HYL1 were used for normalization, respectively.

Given that miRNAs acting in non-cell-autonomous pathways are preferentially processed co-transcriptionally (12), the reduced levels of this type of processing in *hst* mutants may explain the reduced miRNA movement observed in these plants, as well as the recovery in the double mutants. Still, it remains unclear how HST, and co-transcriptional processing of pri-miRNAs, affect miRNA movement. The interaction of HWS with HST and the nuclear pore suggests that transcription and co-transcriptional processing of pri-miRNAs may occur at the nuclear pore in an HWS/HST-dependent manner, perhaps promoting the shuffling of AGO1-unloaded miRNAs out of the nucleus.

To test this hypothesis, we first conducted ChIP-qPCR analysis of *MIRNA* loci associated with HWS, revealing that this protein, as we previously observed for HST (10), interacts with the tested *MIRNA* loci (Figure 5B). Interestingly, this interaction was drastically reduced in *hst-15* mutants suggesting that the *HWS-MIRNA* interaction is mediated by HST (Figure 5B). Like HWS, we also found that NUP1 interacts with *MIRNA* loci as revealed by ChIP-qPCR (Figure 5C). Furthermore, we found that NUP1 also interacts with pri-miRNAs as shown by RIP-qPCR experiments (Figure 5D). This supports the idea that the nuclear pore is not only associated with *MIRNA* loci but that these loci are actively transcribed on the spot rather than being transcribed in the nucleoplasm and relocated to the nuclear pore for export or processing.

Based on this evidence, it is likely that the nuclear pore serves as an active transcriptional site for miRNA loci, a function that was previously shown for other coding genes (29,45). To investigate whether miRNA transcription occurs associated with the nuclear envelope and whether this depends on HST, we used fluorescence in situ hybridization (FISH) to visualize pri-miRNAs within the nucleus using confocal microscopy. We selected *MIR156A,* which produces pri-miRNAs that are processed both at co-transcriptional and post-transcriptional level and contains introns that allow us to differentiate nascent pri-miRNAs from mature transcripts. Using specific FISH probes, pri-miRNA active transcription sites can be then detected as discrete nuclear speckles (one or two per cell) (12). We then measured the distance of these speckles from the nuclear periphery, as previously described (29). In wild-type plants, most pri-miRNAs were located near the nuclear envelope, with a moderate bias toward the nucleoplasm in miRNA biogenesis mutants, particularly in *hyl1-2* mutants, which are known to interact with miRNA loci at the chromatin (23,46,47) (Figure 5E). In wild-type cells with two pri-miRNA speckles, we often observed one speckle near the periphery and the other in a more interior location (Figure 5E). When we tested pri-miRNA transcription sites in *HST* and *HWS* mutants, we observed a lower peripheral distribution in *hst* mutants when compared to *hws*. However, consistent with the suppression of the *hst-15* phenotype by *hws-35,* the reduced peripheral distribution observed in *hst* mutants was restored in *hst-15/hws-35* double mutants (Figure 5F). These data suggest that pri-miRNAs, at least some of them, are transcribed and likely processed at the nuclear pore, with HST facilitating the positioning of *MIRNA* loci at the NPC. Conversely, HWS appears to negatively regulate this process, as its mutations restore the peripheral distribution of pri-miRNAs even in the absence of HST, possibly allowing for a cryptic redundancy between HST and other exportins. Supporting the scenario in which HWS negatively regulates pri-miRNA transcription at the NPC, FISH experiments in plants overexpressing *HWS*, but not HWS^ΔFBox^ revealed an internal distribution of pri-miRNAs within the nucleus (Figure 5G).

These results highlight an opposing effect of HST and HWS on pri-miRNA positioning within the nucleus. While HST’s role in this regard may be attributed to its function as an exportin facilitating the movement of *MIRNA* loci toward the NPC, the mechanisms by which HWS opposes this process are less intuitive. A clue to this mechanism may come from the HWS mass-spectrometry experiments showing its interaction with nine MEDIATOR proteins. Among them, the interaction between HWS and MED37, the same subunit we previously found to interact with HST (10), was particularly strong. We further confirmed the interaction between HWS and MED37 by Y2H and BiFC (Figure 5H and I). Although the interaction was detected with the full-length HWS, it was particularly strong when we used the HWS^ΔFBox^ mutated version (Figure 5H and I). This suggests that HWS may induce MED37 degradation, reducing its capacity for interaction. In agreement with this, the overexpression of HWS, but not HWS^ΔFBox^, caused a decrease in tagged versions of MED37D and MED37E, as detected by Western blot and fluorescence quantification (Figure 5J and 5K). This reduction was even more pronounced when we focused on MED37 associated with miRNA loci, as detected by Western blot after DCL1/HYL1 immunoprecipitation (Figure 5L).

These results suggest that HST, through its karyopherin nature, may help recruiting *MIRNA* loci to the nuclear pore by interacting with the MEDIATOR complex. This allows miRNA transcription and likely processing to occur attached to the pore facilitating a more efficiently export of miRNAs due to their proximity with the nuclear pore. Interestingly this process is disrupted by HWS by affecting MED37 turnover. This could lead miRNA transcription and processing to occur in the nucleoplasm switching the way miRNAs are loaded into AGO1, their nuclear export, and ultimately their cell-to-cell movement.

## Discussion

For many years, HST was believed to play a singular and specific role in miRNA biogenesis: the transport of mature miRNAs from the nucleus to the cytoplasm (2,47). However, recent studies have revealed additional functions of HST in this pathway (10–12). For example, we found that HST acts during the early stages of miRNA biogenesis, attracting DCL1 to *MIRNA* loci through its interaction with the MEDIATOR Complex (10). This action promotes the co-transcriptional processing of pri-miRNAs (12). Furthermore, HST appears to be critical to allow non-cell-autonomous movement of miRNAs (11), although the exact mechanism behind this function remains unclear. A link between the above-described roles of HST in co-transcriptional miRNA processing and miRNA movement is also evident by the fact that mobile miRNAs are predominantly processed in a co-transcriptional manner (12), suggesting that the mobile nature of miRNA may be defined by their processing mechanism (48,49).

Through a suppressor screen of *hst-15*, we identified a genetic interaction between HWS and HST in the miRNA biogenesis pathway, an interaction that although previously described remains poorly understood (37). The nuclear pore complex (NPC) is a highly conserved multiprotein structure composed of a cytoplasmic ring, a central canal, and two nuclear rings connected by eight filaments. The nuclear basket, part of this structure, face the nuclear side of the pore and plays key regulatory roles in gene expression by remodeling chromatin around the NPC, a process dependent on NUA, orthologous to TPR in yeast (4,27,28,45,50,51). We found that HWS interacts with NUA and NUP1, two nuclear basket proteins on the inner side of the nucleus, thereby localizing HWS to the NPC. Interestingly, NUP1 has previously been linked to miRNA biogenesis through its interaction with the TREX-2 complex, which mediates miRNA transcription and export (17). Another protein associated with the NPC is THO2, which facilitates mRNA transport and is also involved in miRNA biogenesis. Mutants of THO2 show reduced miRNA levels, although it does not interact with any component of the pri-miRNA processing machinery (25). Here we show that HWS also interacts with proteins involved in nuclear-cytoplasmic trafficking, such as exportins, importins, RAN proteins, and WIP proteins. Our findings suggest that transcription and pri-miRNA processing occur near or in association with the NPC. HST, due to its interaction with the MEDIATOR complex, its exportin nature, and its role in co-transcriptional pri-miRNA processing (7,10,12) could facilitate the relocation of *MIRNA* loci to the nuclear pore where they are transcribed and processed allowing a more efficient shuttling of mature miRNAs out of the nucleus (Figure 6). The interaction of HWS with many karyopherins, the fact that the mutation of *HWS* partially suppresses *hst* phenotype, and that its overexpression exacerbates it, suggest that in the absence of HST, other exportins may compensate, although less efficiently, for its function. In this sense, in the absence of HWS some of these exportins would escape this proteińs negative regulatory effect partially compensating *hst* mutant phenotype. In the same line of reasoning, the fact that the overexpression of HWS increases *hst* phenotype suggests that in normal conditions HWS acts only in a fraction of the NPC, still allowing some exportin to partially overcome *hst* absence. In this sense, the overexpression of HWS may saturate the NPC preventing any partially redundant effect from happening. This is also evident in our FISH experiments where the peripheral location of *MIRNA* loci, already reduced in *hst* mutants, becomes even lower if HWS is overexpressed (Figure 6).

**Figure 6.**
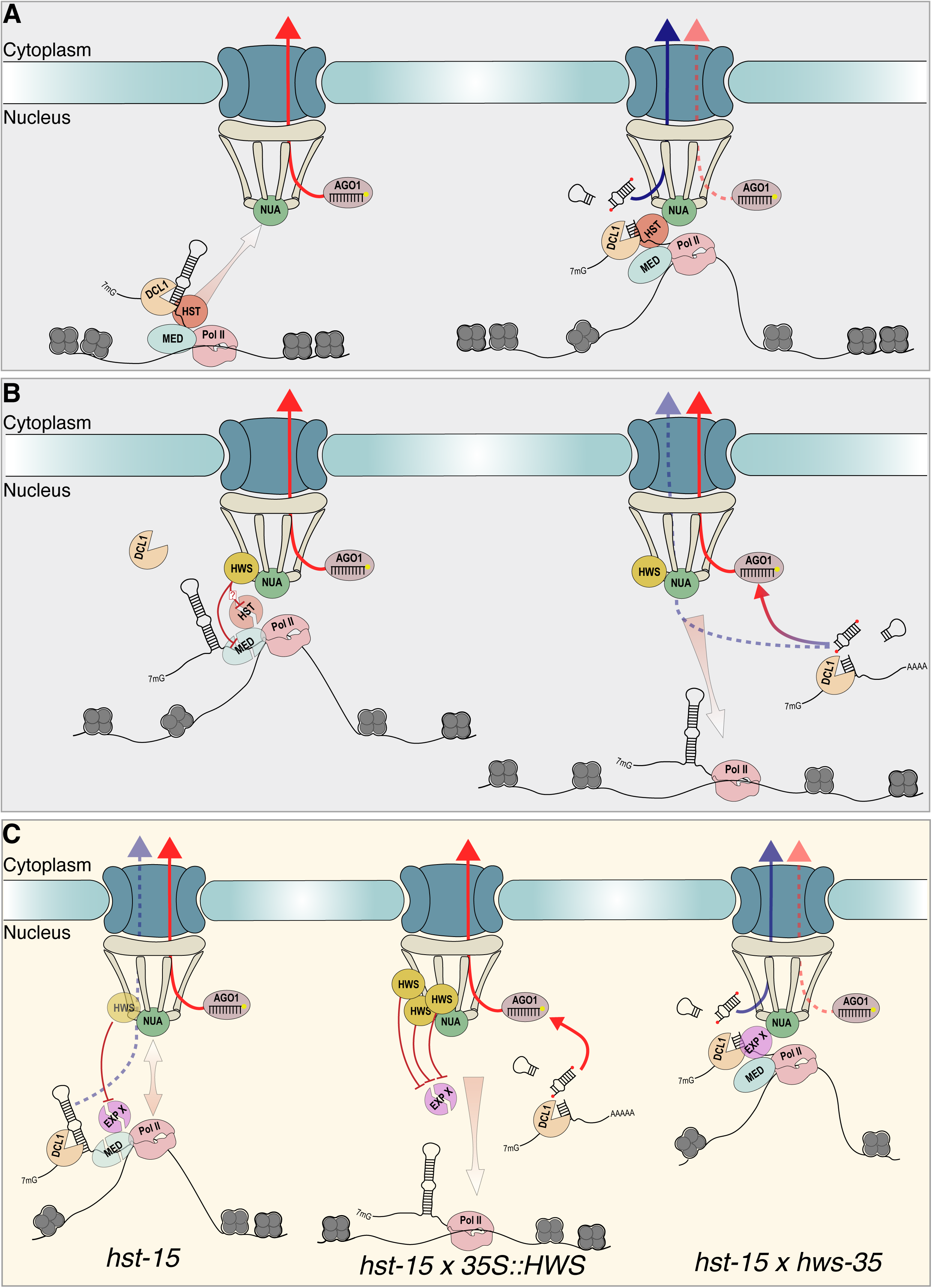
An hypothetical model for NPC-associated miRNA biogenesis and export. A-Given its exportin nature and its capacity to interact with the MEDIATOR and miRNA processing complexes, HST drags *MIRNA* loci to the NPC. RNA Pol II transcription and DCL1-mediated co-transcriptional processing of pri-miRNAs at this location likely facilitate miRNA export from the nucleus in an AGO1-independent manner (blue arrows), promoting non-cell-autonomous functions of mobile miRNAs. Cell-autonomous miRNA shuttling is indicated with red arrows, while blue arrows denote pathways prone to non-cell-autonomous movement. B-HWS interacts with the NPC to antagonize the HST-mediated recruitment of MIRNA. By inducing the degradation of MED37, and possibly HST, HWS causes the detachment of *MIRNA* loci from the NPC, leading to nucleoplasmic post-transcriptional pri-miRNA processing, which, due to its nuclear association with AGO1, restricted miRNAs to cell-autonomous functions. C-In the absence of HST non-cell-autonomous miRNA movement is severely impaired causing the observed mutant phenotype. It is likely that a partially redundant exportin, though inefficient in the miRNA pathway, may still act at NPCs not associated with HWS. This is supported by the fact that the *hst* phenotype is even more pronounced when HWS is overexpressed, likely preventing inefficient exportins from partially compensating for HST. Conversely, in the absence of both *hst* and *hws,* this weak redundancy appeared to be sufficient to partially rescue *hst* phenotype, restoring miRNA cell-to-cell movement. Gray background upper boxes (A and B) represent the hypothetical interaction between miRNA components and the NPC in wild-type cell at HWS-free (A) or HWS-occupied (B) NPCs. Yellow background lower box (C) represents hypothetical crosstalk between the miRNA pathway and the NPC in different mutants.

HWS, as an F-BOX protein, interacts with components of the SCF complex (33,35) and is expected to target proteins for ubiquitin-mediated degradation. While our results indicate that HWS is not involved in HST degradation, we did find that HWS targets and degrades a partner of HST, specifically a subunit of the MEDIATOR complex known to help HST anchor to *MIRNA* loci (10). It is important to highlight that we cannot fully discard the possibility that HWS also degrades a specialized fraction of HST, for example, the portion attached to the nuclear pore. This is particularly true, given that our experiment to test HWS-mediated HST degradation were done with plants overexpressing a tagged version of HST, which might hide any fraction-specific effects on HST turnover. In any case, the degradation of MED37, seems to lead to the detaching of HST from *MIRNA* loci explaining why *MIRNA* loci are no longer located at the nucleus periphery when HWS, but not HWS^ΔFBox^, are overexpressed according to our FISH experiments.

Such detachment of the *MIRNA* loci from the NPC is not expected to affect miRNA biogenesis, which correlates with the largely unaffected levels of miRNAs in *hst* or *hws* mutants (11,35). However, it would affect the subcellular location of miRNA processing, likely impacting how pri-miRNAs are processed, whether and where miRNAs are loaded into AGO1, and the efficiency of the nuclear transit of miRNAs.

It has been reported that the *hst* phenotype arises from defects in miRNA movement rather than from a significant reduction in miRNA levels (11,35). Our findings in the *hst/hws* double mutant, where miRNA movement is restored suggest that HWS influences miRNA function rather than its synthesis. In *hws* mutants, miRNA loading into AGO1 is impaired, which could explain the restored mobility of miRNAs in the double mutant as AGO1-unloaded miRNAs were proposed as the mobile molecules (14). Interestingly, our mass spectrometry analysis revealed the interaction between HWS, the NPC, and AGO1, but AGO1 also appeared to interact with the NPC (17), which position HWS, AGO1, HST, and the NPC in close communion. Although the general pool of AGO1 does not appear to be affected by HWS (35), HWS might selectively degrade AGO1. This type of selective degradation could target specific fractions of AGO1, for example, the fraction associated with the nuclear pore affecting only specific AGO1 functions. Although it remains to be determined which is the role of the NPC-associated AGO1, it is tempting to hypothesize that its preferential location, in combination with our reported NPC-associated miRNA processing, allows for a quick and efficient loading of mature miRNA directly over the NPC facilitating their export.

Another intriguing finding is that mimicry-attenuated miRNA activity, including mimicry-induced miRNA degradation, no longer functions properly in *hws* mutants (35,36). This observation may also point to HWS degrading a specific fraction of AGO1, in this case the pool of AGO1 associated with spurious targets, a fraction of AGO1 prone to degradation (52–54). In this case, it is also possible that other F-BOX proteins, such as FBW2 known to induce AGO1 degradation (52,55), act redundantly or cooperatively with HWS. Whether this level of regulation exists, and whether it is also linked to the nuclear pore, remains to be addressed.

Given that *hws* mutants only show mild phenotypes, it is likely that HWS’s functions described here may not be uniform across all NPCs and tissues. This suggests that HWS-mediated regulation likely takes place at specific NPCs and in certain cell types or tissues. Still, based on our data indicating that HWS negatively regulates co-transcriptional miRNA processing and movement, its absence is not expected to cause a general effect on miRNA biogenesis. HWS’s broader localization in the cytoplasm and nucleus also implies that it may be involved in additional processes beyond its role in miRNA biogenesis.

Our findings offer a new perspective on HST and HWS’s roles in miRNA biogenesis and function but, most importantly, position the NPC as a central hub controlling miRNA processing and the fate of mature miRNAs. The interplay between HWS, HST, and the nuclear pore may provide a mechanism for miRNA movement, AGO1 loading, and regulation of miRNA activity (Figure 6). However, further research is needed to clarify the roles of these proteins, particularly in relation to their localization and selective degradation functions, which may vary across different tissues and individual NPCs.

## Material and Methods

### Plant Material and Growth Conditions

We used *A. thaliana* Col-0 ecotype and the mutant lines *hst-15* (SALK_079290), *hst-6* (2), and *hws-1* and *hws-3* (*35*). The double mutants *hst-15/hws-35*, *hst-15/hws-1*, *hst-15/hws-3*, *hst-6/hws-35*, *hst-15/hws-1,* and *hst-6/hws-3* were generated for this study. Mutations were confirmed by PCR on genomic DNA using specific oligonucleotides (see Table S2). Seeds were disinfected with 70% (v/v) ethanol for 5 min, followed by 30 min in 10% (v/v) sodium hypochlorite and 0.1% (w/v) SDS, then washed with sterile water, and stratified in darkness at 4 °C for 2-3 days. Seedlings were grown on Petri dishes containing semi-solid Murashige and Skoog medium (Sigma-Aldrich) 0.5X (pH 5.7) with 0.6% (w/v) agar or in pots with substrate depending on the experiment. Plasmids used to transform *N. benthamiana* or *A. thaliana* are listed in Table S3. 35S::mCitrineHWS, 35S::mCitrine:HWS^ΔFBox^, proADH1:GAL4AD:HWS and proADH1::GAL4AD:HWS^ΔFBox^ were previously reported (35). Primers used for cloning are listed in Table S2. Genes were amplified using Phusion polymerase (Thermo Fisher), cloned in pEntr/D-TOPO (Thermo Fisher), and recombined using LR Clonase (Thermo Fisher) into Gateway-compatible pGreen destination vectors.

### EMS Mutagenesis, Suppressor Screening, and Mutation Mapping

To perform seed mutagenesis of *hst-15* with the ethyl methane sulfonate (EMS), 300-400 mg of seeds were aliquoted and hydrated with a 0.005% (v/v) Tween-20 solution for 2-4 hours. Seeds were washed with water and mixed with 0.15% (v/v) EMS, incubating for 16 hours with rotation. Eight washes with water were performed for 20 min each with continuous rotation. In the final wash, seeds were resuspended in 0.1% (w/v) agar and stratified for 3 days at 4°C in darkness. Each pot was harvested separately, resulting in approximately 300 groups of about 60 plants each. These groups were used for suppressor screening, with mutant plants of interest identified by their phenotypes. Aberrant plants or those showing phenotypes similar to *hst-15* were discarded. Selected double mutant plants were backcrossed with parental line plants to generate mapping groups for whole-genome sequencing. Mapping groups were composed of samples taken from approximately 300 plants for each mutant, and genomic DNA extracted using the CTAB method. Libraries were prepared with an in-house scaled-down version of Illumina’s TruSeq protocol (10), and sequenced on an Illumina HiSeq3000 instrument (Illumina, San Diego, CA, USA).

Paired-end sequencing of 150-base reads was performed, yielding an average of 72 million read pairs per set and 25 million for the parental line. Bioinformatic analysis began with the removal of low-quality bases from the 3’ end of the reads using the trimmomatic program (56), resulting in an average read length of 130 bases. Reads were mapped to the *A. thaliana* genome using Bowtie2 (57), with mapping percentages exceeding 90% of the reads. Mappings from multiple lanes were combined, and genomic variants were determined using Samtools and Bcftools (version 0.1.19 compatible with SHORE, (58)). The variants were compared with those present in the *hst-15* line using SHOREmap backcross (version 3.6, (30,59)) with a minimum q-score of 25, and variants with an allelic frequency greater than 0.9 were selected. Finally, variants were annotated using SHOREmap to identify affected genes and potential mutation effects.

### RNA Analysis

Total RNA was extracted using TRIzol reagent (Thermo Fisher Scientific). RNA blots were performed as previously described (60). For quantitative RT-qPCR, 1 μg of total RNA was treated with DNase I (Thermo Fisher Scientific), and cDNA was synthesized using the RevertAid RT Reverse Transcription Kit (Thermo Fisher Scientific). qPCRs were performed on three biological replicates, with Actin 2/8 (At3g18780/At1g49240) used as a reference gene. Fold changes were calculated using the 2^-ΔΔCt^ method. Error is expressed as the standard error of the mean (SEM). Statistical differences (P < 0.05 or P < 0.01) were calculated using ANOVA followed by Tukey’s multiple comparison test or by two-tailed unpaired t-tests with false discovery rate (FDR) corrections for multiple paired comparisons.

### Transient Transformation of *N. benthamiana*

Transient transformation of leaves was performed following a previously described protocol (61). *Agrobacterium tumefaciens* bacteria transformed with constructs of interest were grown in YEB medium at 28 °C for 24 to 36 hours. Cells were scraped and resuspended in 500 μL of wash solution (10 mM MgCl2, 100 mM acetosyringone), then briefly shaken and diluted to an OD600 of 0.5 in infiltration solution (1/4 MS [pH = 6.0], 1% sucrose, 100 mM acetosyringone, 0.005% Silwet L-77 [v/v, 50 ml/l]). Depending on the experiment, transformations were performed with a construct expressing the viral silencing suppressor p19. Bacteria were infiltrated into the abaxial side of tobacco leaves, and plants were kept under light for 1 hour and in darkness for 24 hours, before transferred to the growth chamber for 2-3 days before sample collection/microscopy.

### Confocal Microscopy

For localization, co-localization, and protein-protein interaction, a Leica TCS SP8 confocal microscope was used for visualization and image capture. Images were captured using a 40x lens, exciting samples with lasers at 488 nm and 552 nm, and collecting emissions at 500-530 nm, 600-630 nm, and 500-530 nm for eGFP, mCherry, and mCitrine, respectively. Image processing was carried out using Fiji software (62). In FISH experiments, nuclei were observed under Olympus F3000 confocal microscope using lasers emitting light at wavelengths of 405 (Hoechst) and 488 (Alexa 488). To minimize bleed-through between fluorescence channels, the low laser power (0.4–5% of maximum power) and single-channel collection were applied.

### Root Staining with Basic Fuchsin

The staining was carried out using an adaptation of a previously described protocol (11). Five-day-old seedlings grown in vitro on vertical plates in MS medium were used. Roots were cleared in 1 M KOH for 3 hours at 37°C, washed once with distilled water, stained with 0.01% basic fuchsin for 10 minutes with gentle agitation, washed twice in 70% ethanol, and rehydrated in water before imaging with a confocal microscope.

### Separation of Vascular and Epidermal Tissues from Leaves

To separate vasculature from the epidermis, 18-day-old post-germination *A. thaliana* plants were used, and rosette leaves were placed in a paper tape “sandwich” following previously published protocol (38). This method allowed for the separation of the lower epidermis from the vasculature and other tissues present on the upper side of the leaf. Both parts were processed separately RNA extraction.

### Protein Blots

Approximately 300 mg of 15-day-old *A. thaliana* seedlings or three leaves of *N. benthamiana* were processed by grinding in liquid nitrogen. Subsequently, 100 μl of protein extraction solution (50 mM Tris pH 7.5, 150 mM NaCl, 1 mM EDTA, 10% (v/v) glycerol, 1 mM DTT, and one tablet of Complete Protease Inhibitor Cocktail [Roche]) was added. The mixture was centrifuged for 20 min at 16,000 g and 4°C. Proteins were separated on SDS-PAGE (8%) gels and transferred to a PVDF membrane (Amersham). The membrane was incubated with the appropriate primary (anti-GFP (Agrisera AS152987); anti-HYL1 (Agrisera AS09532); anti-ACTIN8 (Agrisera AS132640)) and secondary (polyclonal IgG conjugated with HRP (Agrisera AS09602)), antibodies to detect proteins of interest. The membrane was then exposed on ECL Hyperfilm radiographic films (Amersham) at room temperature for varying times depending on the desired signal intensity.

### Protein-Protein Interaction

For BiFC assays, *N. benthamiana* plants were transformed as previously described, and fluorescence intensity in each sample was detected by confocal microscopy. Yeast Two-Hybrid (Y2H) assays were carried out following the “Yeast Protocols Handbook” (Clontech Laboratories). Competent cells were transformed with 1 μg of DNA from each construct. After electroporation, *S. cerevisiae* cells were recovered in 1 mL of 1 M sorbitol, incubated at room temperature for 5 minutes, and plated on selective YNB-LT medium (7.6 g/L Yeast Nitrogen Base [YNB], 20 g/L glucose, 0.64 g/L DO supplement -leu -trp (Clontech), 15 g/L agar) supplemented with 1 M sorbitol. Two to three yeast colonies co-transformed with the constructs were resuspended in sterile distilled water, adjusted to an OD600 = 0.5, and subjected to three serial 1/10 dilutions. The dilutions were plated on selective -LT medium as a growth control and on -LTH medium (7.6 g/L Yeast Nitrogen Base [YNB], 20 g/L glucose, 0.62 g/L DO supplement -leu -trp -his [Clontech], 15 g/L agar) for the protein interaction assay. Plates were incubated at 30°C for 3-5 days until colony growth was observed. The -LTH selection medium was supplemented with 10 mM 3-AT (3-amino-1,2,4-triazole) to reduce autoactivation by some constructs.

### Co-Immunoprecipitation for Analyzing Protein Degradation

Approximately 3 g of plant material was crosslinked using 37% formaldehyde and then frozen in liquid nitrogen. First, nuclei extraction was performed by resuspending pulverized material in Extraction Solution I (10 mM Tris-HCl pH 8, 0.4 M sucrose, 10 mM MgCl2, 5 mM β-mercaptoethanol, 200 μM PMSF). The homogenate was filtered through a Miracloth layer, and the aqueous phase was centrifuged at 1,500 g for 10 min. The pellet was washed twice with Extraction Solution II (10 mM Tris-HCl pH 8, 0.25 M sucrose, 10 mM MgCl2, 5 mM β-mercaptoethanol, 1% Triton X100, 200 μM PMSF), and then resuspended in 500 μl of Extraction Solution III (Tris-HCl 10 mM pH 8, 1.7 M sucrose, 2 mM MgCl2, 5 mM β-mercaptoethanol, 0.15% Triton X100, 200 μM PMSF). The suspension was carefully pipetted onto 700 μl of Extraction Solution III. The sucrose gradient obtained was centrifuged at 16,000g and 4°C for 5 min. For protein extraction, the pellet was resuspended in 100 μL of lysis buffer (Tris-HCl 50 mM pH 8, 1% NP40, 10 mM EDTA, 200 μM PMSF) and sonicated in a Bioruptor Pico sonicator (Diagenode) (5 cycles of 30 s pulses at high intensity/30 s rest). The mixture was centrifuged at 16,000 g for 10 min at 4°C, and the supernatant was collected. Immunoprecipitation was performed using Sure Beads™ Protein-A magnetic beads (Bio-Rad) following the manufactureŕs instructions. After incubation with the magnetic beads, washes were performed with Chip dilution buffer (1.1% Triton X100, 1.2 mM EDTA, 16.7 mM Tris-HCl pH 8.0, 120 mM NaCl), and samples were incubated for 1 hour at 65°C to reverse the crosslinking. Then, 200 μL of Laemmli sample buffer with added urea (0.24 g/mL) was added, and the samples were incubated for 10 minutes at 95°C. Proteins interactions were detected by Western blot using the corresponding antibodies.

### Nuclear-Cytoplasmic Fractionation

Nuclear-cytoplasmic fractions were performed as described (63) with minor modifications. In brief, 3 g of 2-week-old seedlings were harvested and ground in 6 mL of lysis buffer (20 mM Tris-HCl [pH 7.5], 20 mM KCl, 2 mM EDTA, 2.5 mM MgCl2, 25% glycerol, 250 mM sucrose, and 5 mM DTT). The suspension was filtered through a double layer of Miracloth (Calbiochem) and centrifuged at 1,500 g for 10 min. The supernatant was then centrifuged at 10,000 g for 10 min at 4 °C and collected as the cytoplasmic fraction. The pellet was washed four times with 10 mL of nuclear resuspension buffer NRBT (20 mM Tris-HCl [pH 7.4], 25% glycerol, 2.5 mM MgCl2, and 0.2% Triton X-100), then resuspended with 500 μl of NRB2 (20 mM Tris-HCl [pH 7.5], 0.25 M sucrose, 10 mM MgCl2, 0.5% Triton X-100, and 5 mM Δ-mercaptoethanol). It was carefully overlaid on top of 700 ml of NRB3 (20 mM Tris-HCl [pH 7.5], 1.7 M sucrose, 10 mM MgCl2, 0.5% Triton X-100, and 5 mM Δ -mercaptoethanol) and centrifuged at 16,000 g for 10 min at 4 °C. The nuclear pellet was directly resuspended in TRIzol reagent for RNA extraction.

### RNA Immunoprecipitation

RNA immunoprecipitation experiments were performed following a previously published protocol (64). Two grams of 15-day-old *A. thaliana* seedlings were collected for H3 and AGO1 immunoprecipitation, and 2 g of transformed *N. benthamiana* leaves were crosslinked with formaldehyde for NUP1 immunoprecipitation. Anti-GFP (Agrisea AS152987), anti-AGO1 (Agrisera AS09527), or anti-H3 (Agrisera AS10710) antibodies and SureBeads Protein A magnetic beads (Bio-Rad) were used to immunoprecipitate NUP1:GFP at the NPC, AGO1 at the RISC complex, or chromatin using H3. A negative control was performed using non-transformed plants or anti-IgG (Agrisera AS09605) and compared with Col-0 plants. Reverse crosslinking was performed with proteinase K (QIAGEN). After discarding the beads, the supernatant was used for RNA extraction using TRIzol reagent (Thermo Fisher Scientific).

### Chromatin Immunoprecipitation (ChIP)

ChIP assays were carried out with modifications from (65). 2.5 g of 15-day-old seedlings were cross-linked with formaldehyde. Enriched nuclei were sonicated in a Bioruptor Pico water bath (Diagenode; 30 s on/30 s off pulses at high intensity for 10 cycles using Bioruptor microtubes). Samples were incubated with anti-GFP (Agrisera AS152987) or anti-IgG (Agrisera AS09605, as a negative control) and immunoprecipitated with SureBeads Protein A magnetic beads (Bio-Rad) for 12 h at 4°C. Reverse crosslinking was performed with proteinase K (QIAGEN). Immunoprecipitated DNA was recovered using a mixture of phenol:chloroform:isoamyl alcohol (25:24:1), precipitated in ethanol, and analyzed by qPCR. Untreated sonicated chromatin was processed in parallel as input samples.

### Mass Spectrometry Data Analysis

The mass spectrometry results of proteins interacting with HWS were obtained from (10,35). Controls used were mCitrine fused to the HWS^ΔFBox^ version and plants expressing only GFP. To determine common interactors between HWS and HST, the VennDiagram (66) tool was used. Gene ontology analyses were performed using the functional annotation package for *A. thaliana* (org.At.tair.db, Genome-wide annotation, R package version 3.19.1) and the enrichGO function from the R package clusterProfiler (67). The results were visualized using dot plots with circles representing the categories colored based on their FDR-adjusted p-value and sized proportionally to the number of genes annotated in each category. Phylogenetic trees were generated using the NGPhylogeny server (https://ngphylogeny.fr/, (68)). The processing sequence was as follow: sequence alignment with MAFFT (69), alignment trimming with BMGE (70), and phylogenetic tree inference with PhyML (71), a maximum-likelihood-based program. To assess clade reliability, 100 bootstrap replicates were performed, and nodes with bootstrap values below 50% were removed from the final tree. The trees were visualized using the FigTree program (http://tree.bio.ed.ac.uk/software/figtree/).

### Fluorescent In Situ Hybridization (FISH)

For these experiments, nuclei were isolated from 4-week-old *A. thaliana* leaves fixed for 1 hour in 4% paraformaldehyde in phosphate-buffered saline (PBS), pH 7.2 (72). For pri-miRNA detection, antisense DNA oligonucleotides labeled with digoxigenin at their 5’ ends, recognizing the intron of pri-miRNA156a (Table S2), were applied. Terminal transferase (TdT) (Sigma Merck) was used to add additional digoxigenin-conjugated nucleotides to the 3’ ends of each probe. For this reaction, each probe (at a final concentration of 10 pM) was incubated in reaction buffer (5 mM CoCl2, 0.1 mM DIG-11-dUTP (Sigma Merck), 0.1 mM dATP, 0.2 mM Alexa Fluor 488-5-dUTP (Thermo Fisher)) with 400 U of TdT per reaction (Sigma Merck) for 40 minutes at 37 °C. The pri-miRNAs were localized by applying FISH combined with immunolocalization of digoxigenin attached to the 5’ and 3’ ends of the probes as previously described (73). Before hybridization, the nuclei were treated with PBS containing 0.1% Triton X100. Probes were hybridized in hybridization buffer (30% (v/v) formamide, 4x SSC (600 mM NaCl, 6 μM sodium citrate), 5x Denhardt’s solution (0.1% (w/v) Ficoll 400, 0.1% (w/v) polyvinylpyrrolidone, 0.1% (w/v) BSA, 1 mM EDTA, and 50 mM phosphate buffer, pH 7.2)) in a humid chamber overnight at 26 °C. After washing, a primary anti-DIG antibody from mouse (Sigma Merck) or rabbit (Sigma Merck) (diluted 1:100) was added in PBS containing 0.05% acetylated BSA, and the slides were incubated overnight at 10 °C. The nuclei were washed with PBS and incubated with secondary goat anti-mouse antibodies conjugated with Alexa Fluor 488 (diluted 1:100) (Thermo Fisher) in PBS containing 0.05% acetylated BSA for 2 hours at 37 °C. DNA was stained with Hoechst 33342 (Thermo Fisher) and mounted in ProLong Gold antifade reagent (Life Technologies).

## Acknowledgments

This work was supported by grants from Agencia Nacional de Promoción Científica y Tecnológica (PICT-2020-00757 and PICT-2021-I-A-00452), Universidad Nacional del Litoral (CAI+D 2020), and Spanish Ministry of Science and Innovation (PID2022-137037NB-I00 and CNS2023-145312) to P.A.M. A.L.A. is a member of CONICET; L.G. is a fellow of CONICET. P.A.M is a member of CSIC. We thank Catharina Merchante for critically reading our manuscript.

## Author contribution

L.G and D.G performed most experiments. J.F performed Y2H assays, C.Z performed microscopy and ChIP assays, T.G performed all FISH assays, A.L.A executed the informatic analysis. D.C performed mutant libraries sequencing, L.G, D.G and P.A.M conceived the study; L.G, and P.A.M. wrote the manuscript. A.Z, A.J, and P.A.M secure funds for the project and supervised the experimental procedures.

