## Supplemental information for "The Nuclear Pore Complex acts as a hub for pri-miRNA transcription and processing in plants"

**Supplemental Figures**

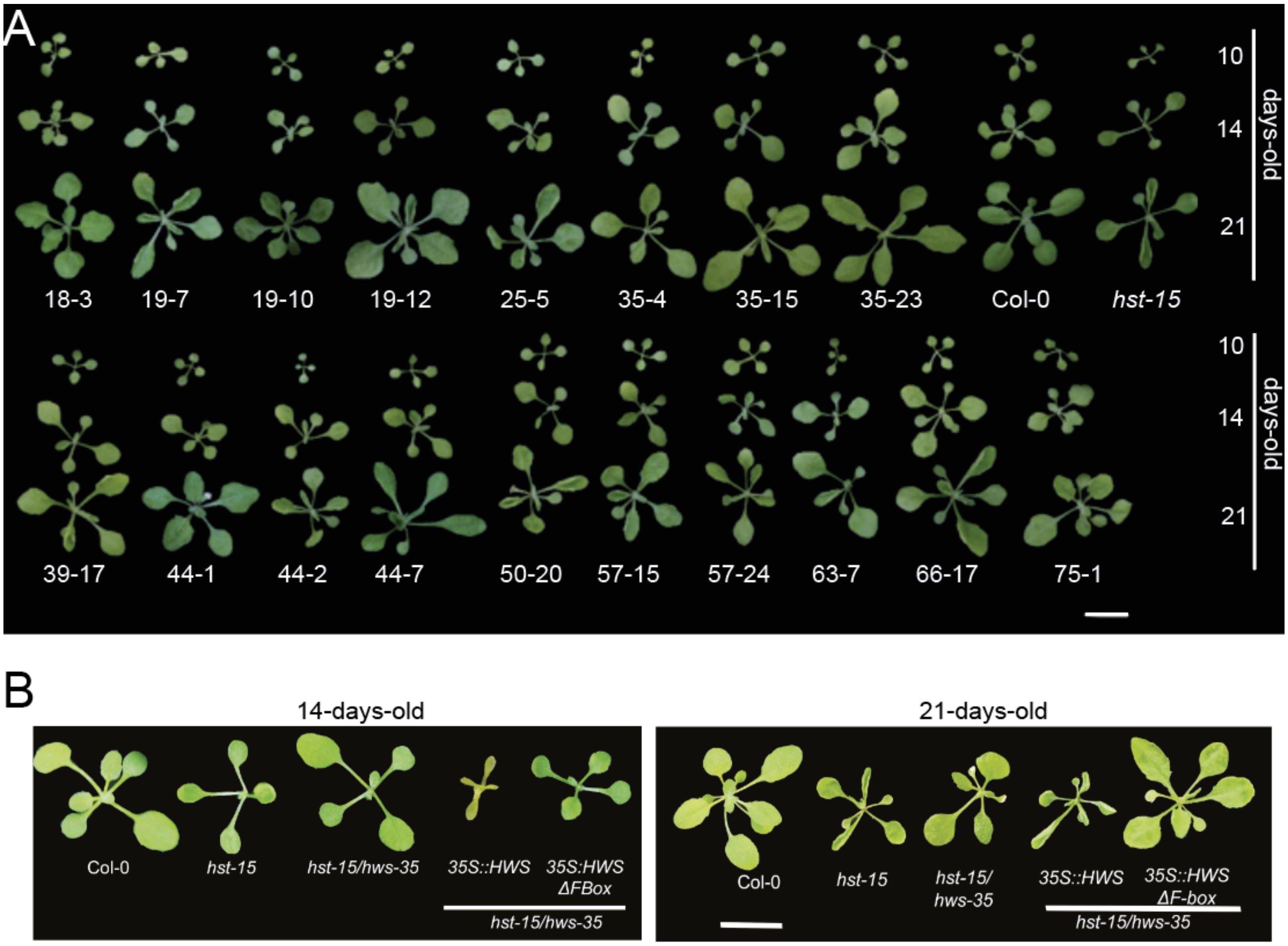


**Figure S1.** Phenotype of 10-, 14-, and 21-day-old mutants isolated from the genetic suppressor screening (A), and complementation lines (B). Scale bar= 1 cm. In all cases the

plants where photographed individually and digitally mounted into a uniform back background to facilitate comparison.

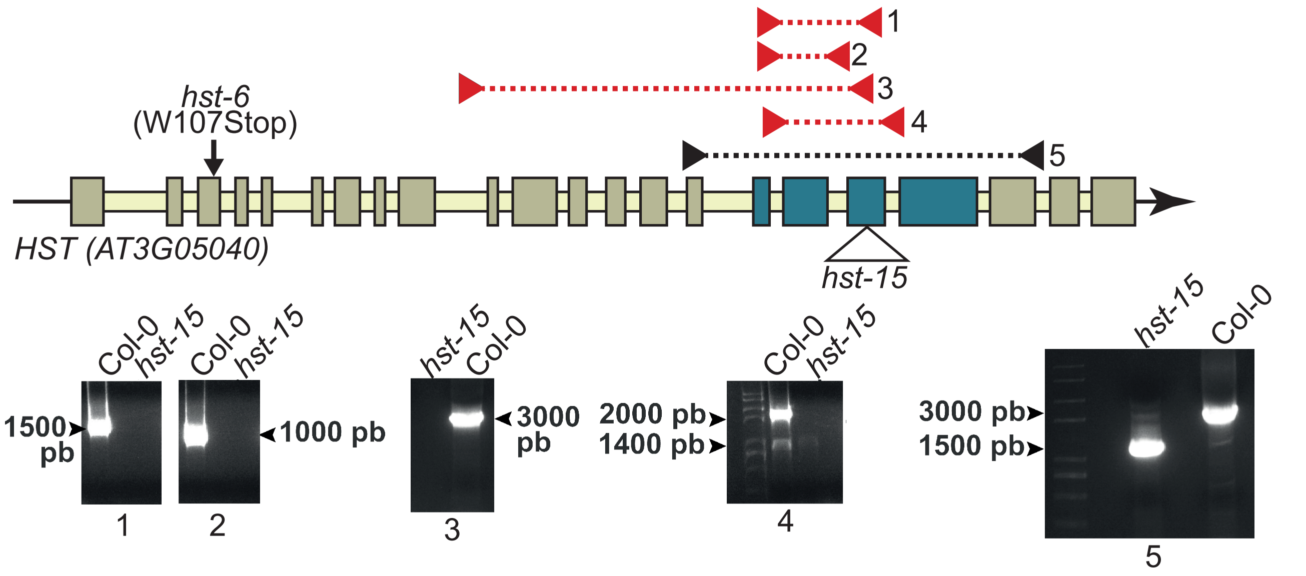


**Figure S2.** Upper panel: Schematic representation of the *HST* gene. Rectangles represent exons, while the thick line refers to introns. Blue rectangles indicate deleted exons due to T-DNA loss. The empty triangle marks the position where the T-DNA was annotated for the *hst-15* allele, and the *hst-6* allele is marked with a black arrow. Red and orange triangles connected by dotted lines represent the non-amplified fragments in *hst-15;* black triangles connected by a dotted line represent the obtained amplification product in this allele.

Lower panel: Photographs of agarose-gel of PCR fragments corresponding to the upper panel-noted amplifications.

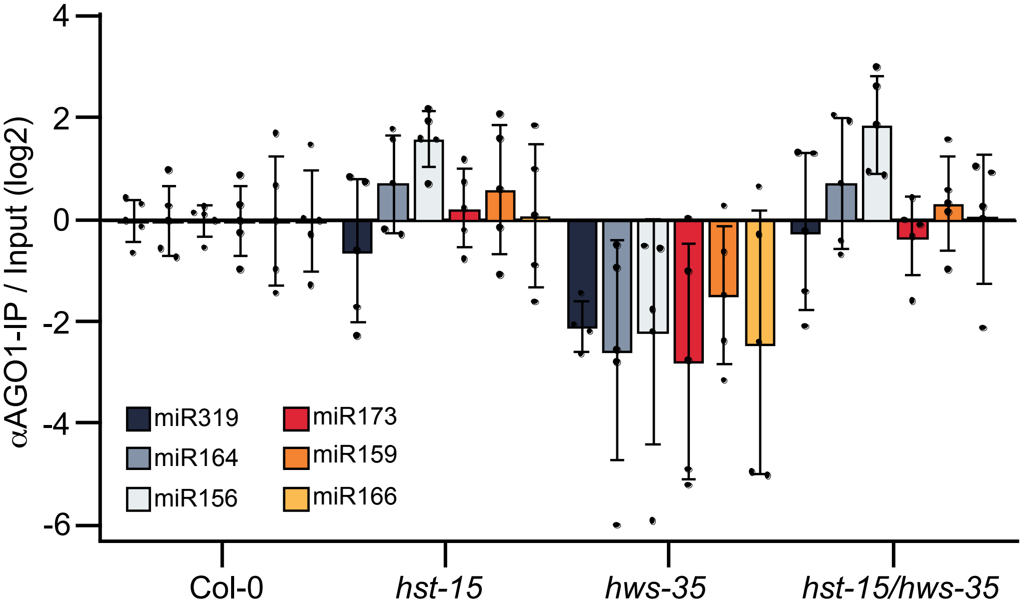


**Figure S3.** RIP-qPCR of miRNAs loaded in AGO1 as detected in IP of total AGO1 in different genotypes. qPCR values are from n=3 biologically independent samples, presented as mean ± SEM. P-values were calculated in a two-tailed, unpaired, t-test.

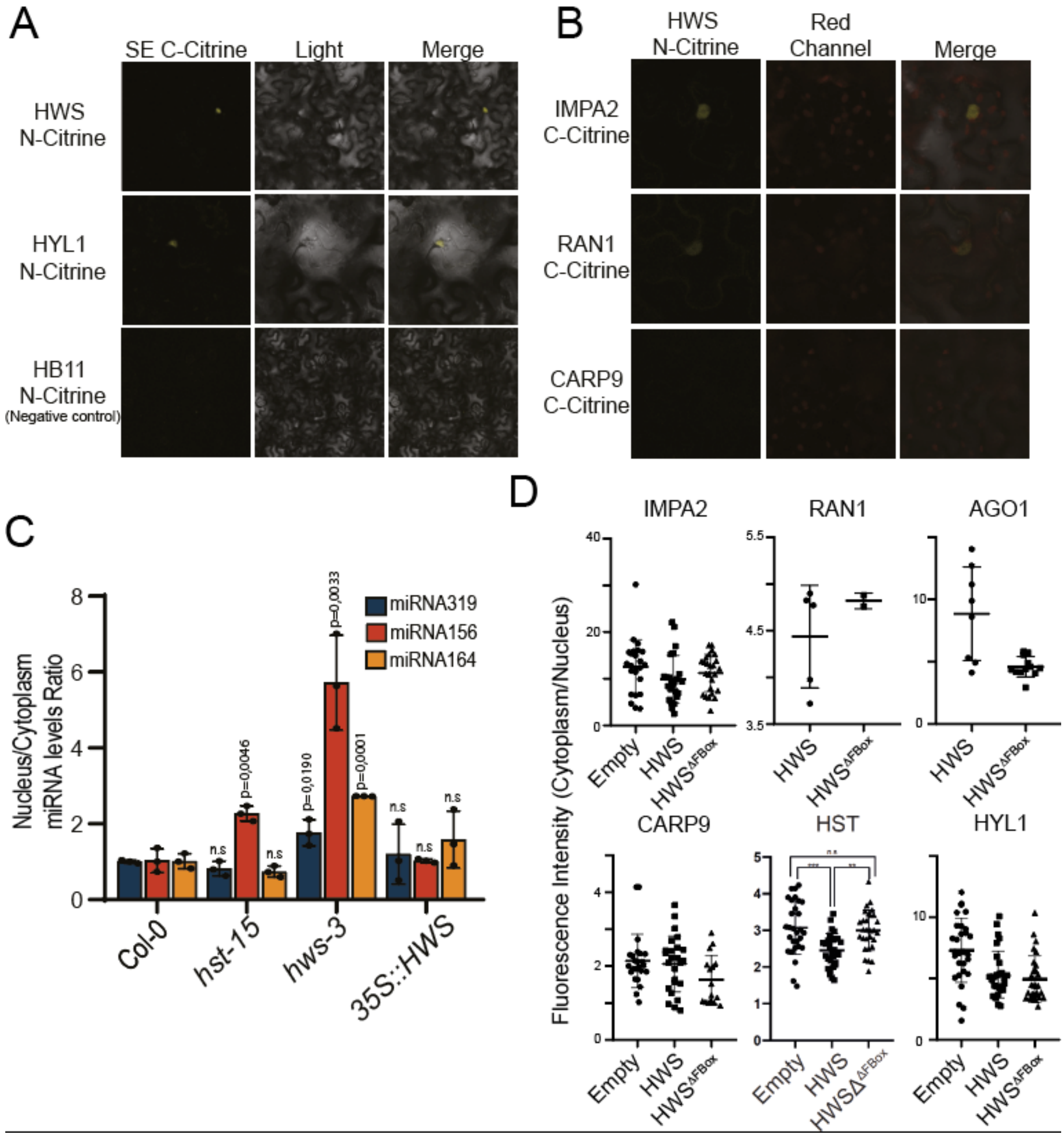


**Figure S4.** A- Interaction between HWS and SE as detected by BiFC, with HYL1 as a positive control and HB11 as a negative control. B- BiFC analysis showing interactions between HWS-IMPA1 and HWS-RAN1, with CARP9 as a negative control. C- Nuclei/cytoplasm fractionation followed by RT-qPCR of miRNAs across different genotypes. D- *N. benthamiana* transformation with IMPA2:GFP, RAN1:GFP, AGO1:GFP, CARP9:GFP, HST:GFP, and

HYL1:GFP, along with 35S:HWS or 35S:HWSΔFBox, to examine differences in subcellular localization.

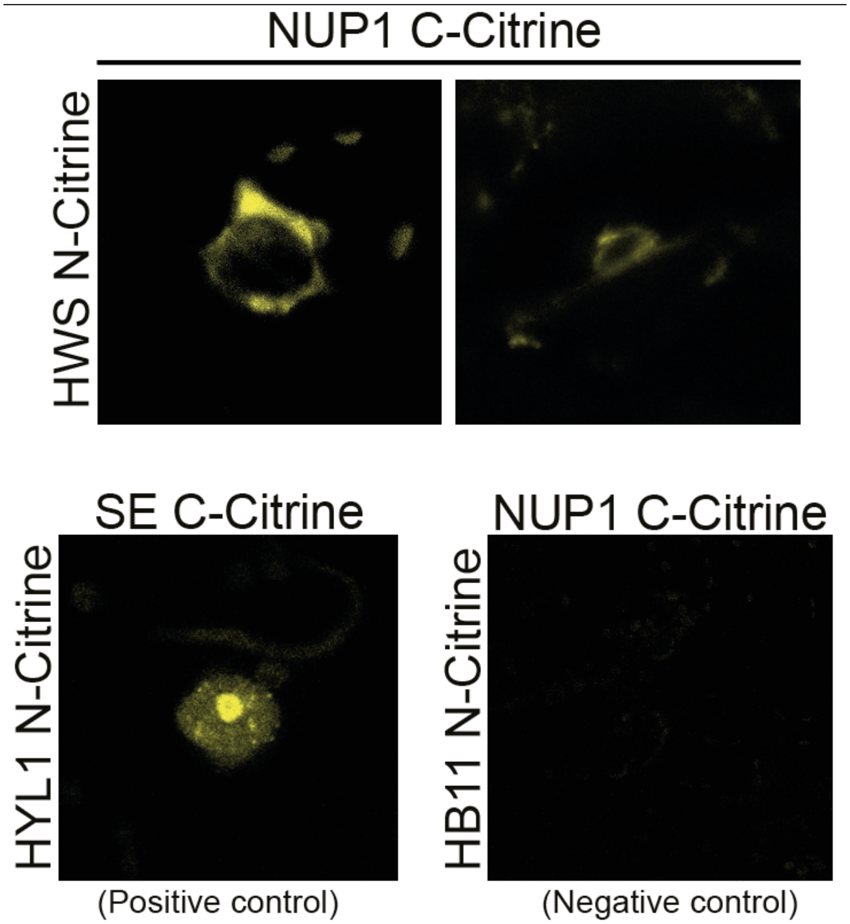


**Figure S5.** Interaction between HWS and NUP1 as detected by BiFC, with HYL1-SE interaction as a positive control and HB11 as a negative control.

**Table S1.** Polymorphisms identified in isolated mutants.

**Table S2.** Common interactors between HWS and HST

**Table S3**. List of primers used in this study.

**Table S4**. List of seeds and clones used in this study.
